# Loss of leaf-out and flowering synchrony under global warming

**DOI:** 10.1101/391714

**Authors:** Constantin M. Zohner, Lidong Mo, Susanne S. Renner

**Affiliations:** Institute of Integrative Biology, ETH Zurich (Swiss Federal Institute of Technology), Universitätsstrasse 16, 8092 Zurich, Switzerland; Systematic Botany and Mycology, Department of Biology, University of Munich (LMU), 80638 Munich, Germany

**Keywords:** Climate change, Phenology, Temperate plants, Flowering, Leaf unfolding, Day-length, Chilling

## Abstract

The temporal overlap of phenological stages, phenological synchrony, crucially influences ecosystem functioning. For flowering, among-individual synchrony influences gene flow. For leaf-out, it affects interactions with herbivores and competing plants. If individuals differ in their reaction to the ongoing change in global climate, this should affect population-level synchrony. Here, we use climate-manipulation experiments, Pan-European long-term (>15 years) observations, and common garden monitoring data on up to 72 woody and herbaceous species to study the effects of increasing temperatures on the extent of within-population leaf-out and flowering synchrony. Warmer temperatures reduce *in situ* leaf-out and flowering synchrony by up to 55%, and experiments on European beech provide a mechanism for how individual genetic differences may explain this finding. The rapid loss of reproductive and vegetative synchrony in European plants predicts changes in their gene flow and trophic interactions, but community-wide consequences remain largely unknown.

---

The structure and functioning of ecosystems crucially depends on the timing of annually repeated life stages, such as leaf-out and flowering (*1–4*). Anthropogenic climate warming is causing advanced leaf-out and flowering in both herbs and trees, and this is affecting growth and reproductive success (*5–8*). Warmer springs and summers are also causing leaf-out and flowering to spread out over longer periods because the sensitivity to winter chilling, spring warming, and day length differs among species (*2,3,9–11*). Such species-specific responses imply variation in heritable phenological strategies among individuals, but how current climate warming is shifting within-population phenology and possibly synchrony has not been addressed. For leaf-out, inter-individual synchrony affects interactions with foliovores and competing plants (*12*). For flowering, reduced inter-individual synchrony should adversely affect gene flow by reducing cross-pollination and fruit set (*13*). To detect such possible effects of climate warming on within-population synchrony, a range of herbs and trees, representing different leaf-out and flowering strategies, needs to be studied.

Here, we use a combination of climate-manipulation experiments, common garden monitoring, and long-term Central European *in situ* observations to analyze effects of warming on within-population phenological synchrony. The long-term data were obtained from the Pan European Phenology Project (http://www.pep725.eu, hereafter PEP). The PEP data consisted of 12,536 individual time series (each minimally 15 years long), comprising the leaf-out times of nine dominant tree species and the flowering times of six tree species, four shrubs, and five herbs (see Methods and the distribution of the sites in Fig. 1a, and Figs. S1 and S2). To define populations, we divided the study area into pixels of one-degree resolution (~110 x 85 km) and then calculated leaf-out synchrony (LOS) and flowering synchrony (FLS) in a given year as the standard deviation of leaf-out or flowering date for all individuals within a pixel (note that the data were cleaned to ensure that observed individuals were the same between years; see Methods). For each pixel and each phenological stage (leaf-out or flowering), we determined the preseason as the period 60 days before the average leaf unfolding or flowering date within the respective pixel.

**Figure 1.**
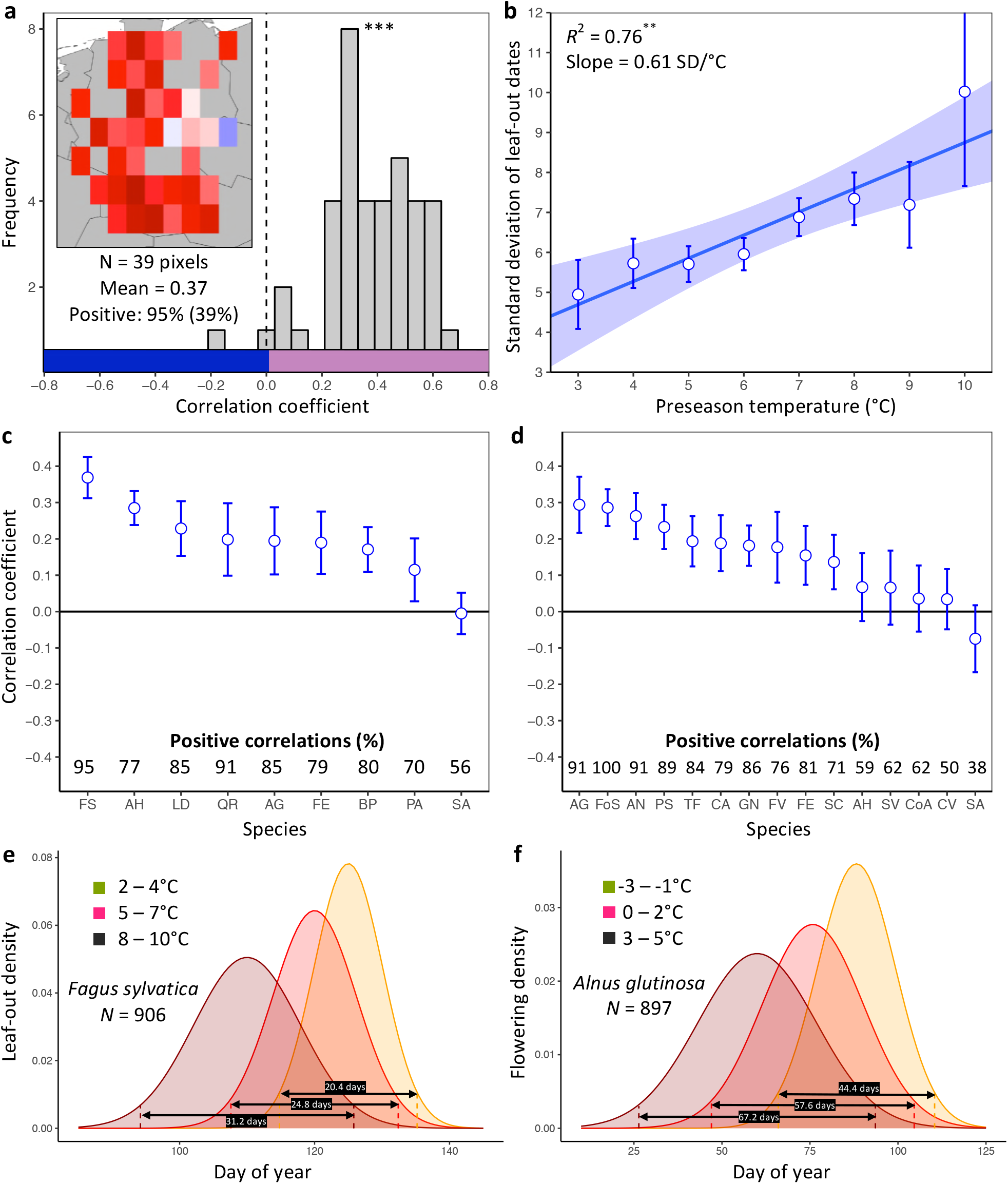
Loss of inter-individual synchrony in leaf-out and flowering with increasing temperatures. **a**, Frequency distribution showing the correlations between the standard deviation of inter-individual leaf-out times and spring temperature for *Fagus sylvatica* at 39 pixels (1° x 1° areas). Mean = Mean correlation coefficients across all sites (*N*), Positive = percentage of positive correlations and the percentage of statistically significant positive correlations (in parentheses). Inset shows a heat map of the correlations at the 39 pixels. **b**, Effect of preseason temperature on the standard deviation of inter-individual leaf-out times (mean ± SEM) in *F. sylvatica* averaged across all years and sites. **c, d**, Mean Pearson correlation coefficients (± 95% confidence intervals) for the effect of spring temperature on the standard deviation of inter-individual leaf-out (**c**) or flowering times (**d**). *Positive correlations* = percentage of the total number of positive correlations. See Figs. S1b and S2b for number of sites (1° x 1° areas) in which the relationship was analyzed. **e, f**, Distributions of inter-individual (**e**) leaf-out dates in *F. sylvatica* and (**f**) flowering dates in *Alnus glutinosa* under different spring temperatures. *N* = Number of available year x pixel (1° x 1° areas) combinations. To model the distributions (means and standard deviations), mixed-effects models were applied including site (pixel) as a random effect. See Figs. S3 and S4 for distributions of all 20 analyzed species. AG, *Alnus glutinosa*; AH, *Aesculus hippocastanum*; AN, *Anemone nemorosa;* BP, *Betula pendula*; CA, *Corylus avellana*; CoA; *Colchicum autumnale*; CV, *Calluna vulgaris*; FE, *Fraxinus excelsior*; FoS, *Forsythia suspensa*; FS, *Fagus sylvatica;* FV, *Fragaria vesca;* GN, *Galanthus nivalis;* LD, *Larix decidua;* PA, *Picea abies;* PS, *Prunus spinosa;* QR, *Quercus robur;* SA, *Sorbus aucuparia;* SC, *Salix caprea;* SV, *Syringa vulgaris;* TF, *Tussilago farfara*.

As expected, within pixels, species’ mean leaf-out dates were negatively correlated with preseason temperature (98% of observation series statistically significant at *P* <0.05), with a mean linear correlation coefficient of −0.76 ± 0.03 (mean ± 95% confidence interval), predicting an average advance of 4.3 ± 0.2 days per each degree warming. Similarly, in more than 99% of pixels, the mean flowering dates were negatively correlated with the preseason temperature (91% statistically significant at *P* <0.05), with a mean linear correlation coefficient of −0.75 ± 0.10, predicting an average advance of 4.6 ± 0.2 days per each degree warming.

Surprisingly, higher preseason temperatures had a negative effect on LOS in eight of the nine species (Figs. 1c and S1) and on FLS in 10 out of 15 species (Figs. 1d and S2). None of the species exhibited a positive effect. Across all species, preseason temperature negatively affected LOS in 78% of analyzed pixels (15% statistically significant at *P* <0.05), i.e., the standard deviation of inter-individual leaf-out times increased by 0.45 ± 0.07 (mean ± CI) days per degree of warming, with a mean linear correlation coefficient of 0.19 ± 0.03. Significant positive effects of preseason temperature on LOS appeared in fewer than 1% of pixels. The species showing the strongest decline in LOS related to warmer preseason temperatures was European beech (*Fagus sylvatica*; Fig. 1a): preseason temperature negatively affected LOS in 95% of analyzed pixels (39% statistically significant), with the standard deviation of inter-individual leaf-out times increasing by 0.61 ± 0.05 days per degree of warming (Fig. 1b) and a mean linear correlation coefficient of 0.37 ± 0.06. When modelling the distribution of within-population leaf-out dates under different preseason temperatures, we found that warming increases the inter-individual variation in leaf-out times by up to 55%, which equates to lengthening the period during which 95% of individuals in a population leaf-out by 11 days (Figs. 1e and S3).

Across all species, preseason temperature negatively affected FLS in 75% of analyzed pixels (18% statistically significant), with the standard deviation of inter-individual flowering times increasing by 0.35 ± 0.15 days per degree of warming and a mean linear correlation coefficient of 0.15 ± 0.06 (Figs. 1d and S2a). A significant positive effect of preseason temperature on FLS was found in only 2% of pixels. The species showing the strongest decline in FLS related to warmer preseason temperatures was the European alder (*Alnus glutinosa*): preseason temperature negatively affected FLS in 91% of analyzed pixels (33% statistically significant), with the standard deviation of inter-individual flowering times increasing by 0.91 ± 0.27 days per degree of warming and a mean linear correlation coefficient of 0.30 ± 0.08. When modelling the distribution of within-population flowering dates under different preseason temperatures, we found that warming increases leaf-out variation by up to 51%, which equates to lengthening the period during which 95% of individuals in a population initiate flowering by 23 days (Figs. 1f and S4). In species, such as the crocus *Colchicum autumnale* and the heath *Calluna vulgaris*, where preseason temperature had little effect on the mean flowering date, preseason temperature also had little effect on FLS (Figs. S4 and S7).

To cross-validate the results obtained from the PEP data, we used common garden data consisting of leaf-out information on 209 individuals in 59 temperate woody species (minimally 3 individuals per species) observed in the Munich Botanical garden from 2013 to 2018. A Bayesian hierarchical model, including preseason temperature as predictor variable, the standard deviation of inter-individual leaf-out times per year as response variable, and species as a random effect, showed a significantly negative effect of preseason temperature on LOS (lower panel Fig. 2a). On average, across all 59 species, the standard deviation of inter-individual leaf-out times increased by 0.26 ± 0.10 (mean ± CI) days per degree of warming.

**Figure 2.**
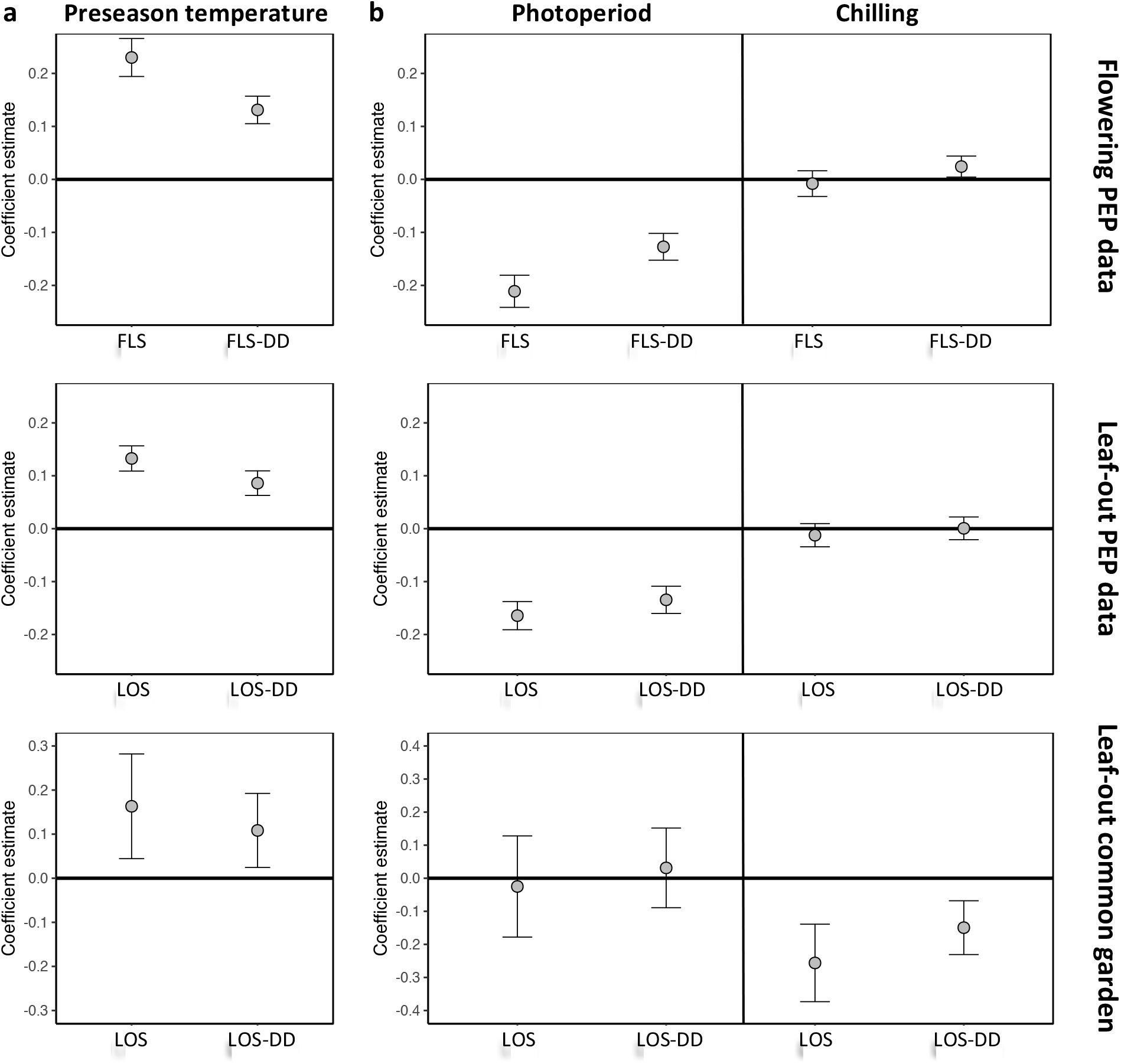
The environmental drivers of inter-individual phenological synchrony as inferred from (i) flowering times (upper panels) and (ii) leaf-out times (middle panels) using the PEP data, and (iii) common garden observations on leaf-out times. **a**, Coefficient estimates (effective posterior means ± 95% credible intervals) for the effect of preseason temperature (mean temperature 2 months before a species’ mean leaf-out/flowering date) on inter-individual phenological synchrony measured either as the standard deviation in leaf-out/flowering dates (LOS / FLS; left) or the standard deviation in degree-day (DD) requirements among individuals (LOS-DD / FLS-DD; right). **b**, Coefficient estimates for the effects of photoperiod and winter chilling on inter-individual leaf-out synchrony. Hierarchical Bayesian linear models were applied using information on 13 (upper), 9 (middle), and 59 species (lower panels). To account for within-species rather than among-species synchrony, all models include species random effects. The models using the PEP data (upper and middle panels) additionally include site random effects (1° pixels) to address within-population phenological synchrony. All variables were standardized to allow for direct effect size comparisons.

Which factors cause the loss of inter-individual synchrony under climate warming? One possibility is that individuals reach their forcing sums required for leaf-out or flowering over a longer period because “within-spring warming speed” may be decreasing, flattening the temperature curve during spring (*14,15*). Thus, while the time span among individual leaf-out times might increase, differences in the forcing sums required until leaf-out or flowering among individuals might remain similar. To test this, we additionally calculated leaf-out/flowering synchrony as the standard deviation in individual forcing requirements (degree-days [DD] from 1 January until leaf-out/flowering) [hereafter referred to as LOS-DD and FLS-DD] for both the PEP and Munich common garden data. In both data sets, we found a strong (albeit slightly weaker compared to the LOS/FLS analysis) negative relationship between preseason temperature and LOS-DD, i.e., individual differences in the forcing sums required until leaf-out or flowering are increasing with warmer preseasons (Figs. 2a and S6). We also simulated synchrony of spring phenology based on the Munich Jan–May temperatures over the past 60 years, assuming that phenology is solely driven by degree-day accumulation (no effect of photoperiod or winter chilling; see Fig. S5b) and this simulation revealed small losses of synchrony (R^2^ values between 0.04 and 0.11 and regression coefficients between 0.15 and 0.43, see Fig. S5c). Together, those results show that a flattening temperature curve during spring can account for only a minor proportion of the declining inter-individual synchrony in the 72 species analyzed here.

Warmer preseasons in spring are associated with both reduced accumulation of winter chilling and shorter day-lengths at spring onset, and previous experiments on plant phenological strategies have shown pronounced differences among species in their reactions to day length and winter chilling (*9–11*). To test whether similar differences within species might explain the decrease in LOS and FLS under climate warming detected in our *in-situ* data, we designed experiments in which we exposed trees to different regimes of spring warming, winter chilling, and day length. We additionally tested for the relative effects of winter chilling and day length on LOS and FLS using the PEP and Munich common garden data (for each year and individual, we calculated the winter chilling experienced until leaf-out and the day length for the date when an individual’s average forcing requirement had been reached).

A first experiment addressed inter-individual variation in spring warming (‘forcing’), day length, and winter chilling requirements in 11 mature *Fagus sylvatica* trees grown in the vicinity of the botanical garden in Munich. Twigs were cut at three dormancy stages during winter and exposed to different day-length regimes (8 h, 12 h, or 16 h light per day) and ambient spring-forcing conditions (mean daily temperature of 16°C). Note that in beech, leaf-out and flowering occur simultaneously because leaves and flowers are located on the same preformed shoots within overwintering buds. The results showed large differences in forcing and day-length requirements among individuals (Fig. 3a and b): for example, while in individual 1, day length had no effect on the amount of warming required until budburst, in individual 11, warming requirements were >2x lower under long-day than under short-day conditions (Fig. 3b). Chilling requirements differed little among individuals (compare slopes in Fig. 3c).

**Figure 3.**
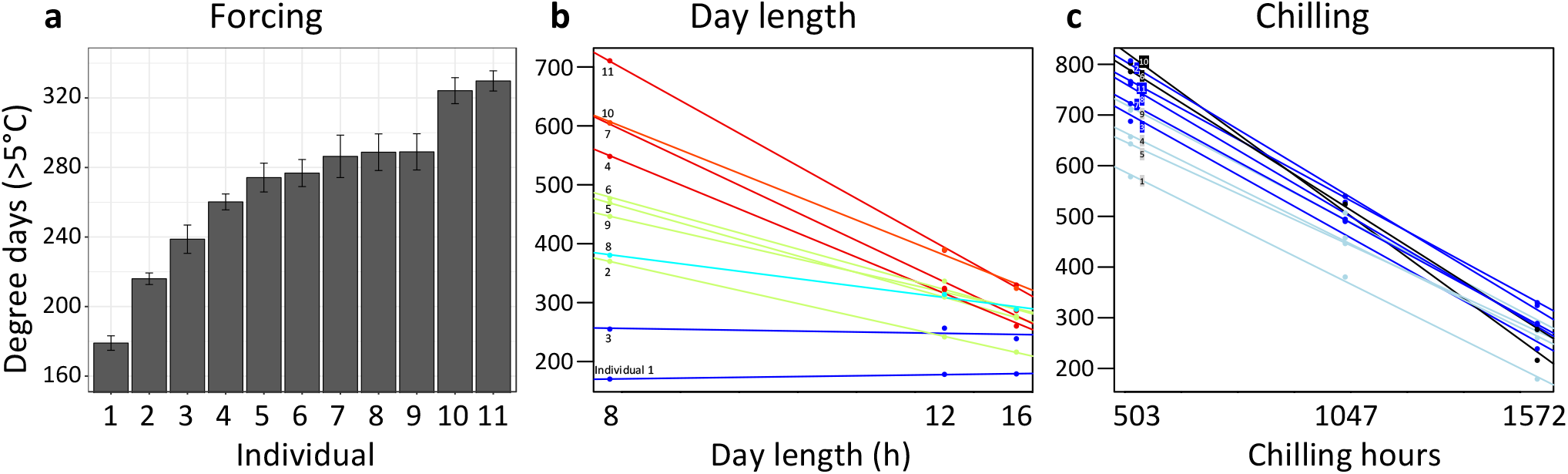
Individual differences in the forcing (a), day-length (b), and chilling (c) requirements among 11 beech trees (*F. sylvatica*; Experiment 1). **a**, Mean (± SEM) forcing requirements (accumulative degree days >5°C) until budburst under long chilling and constant 16-h day length. **b**, Degree days until budburst at 8-h, 12-h, and 16-h day length (collection date: 21 March 2015). Colours according to slope (red: steep slope; blue: no slope). c, Degree days until budburst under short, intermediate, and long chilling (collection dates: 22 Dec 2014, 6 Feb 2015, 21 March 2015) and 16-h day length. Colours according to slope (dark blue: steep slope; light blue: no slope).

In a second experiment, we cut twigs of the same 11 beech trees at eight dormancy stages during winter and exposed them to natural day length. Temperatures were the same as in experiment 1, i.e., ambient. This allowed us to determine (i) the extent to which differential reliance on forcing, photoperiod, and winter chilling (as inferred from experiment 1) explains LOS/FLS under natural light conditions, and (ii) the effect of warmer winter and spring conditions on LOS/FLS. As in the *in situ* data from the Pan European Phenology network, synchrony strongly decreased under warmer spring conditions (Fig. 4 a, b), likely because of day-length sensitivity differences among individuals (as documented for *F. sylvatica*; Fig. 3b): Under cold winter conditions, days are already long when spring warming occurs, reducing the effect of a tree’s day length sensitivity on its leaf-out time, whereas with early spring warming, days are still short, preventing day-length sensitive trees from budburst. In natural populations, leaf-out advancement in day length-sensitive individuals, but not in day length-insensitive individuals, will thus increase the period of leaf-out under short day conditions. Both the experimental and the PEP *in situ* data confirm this idea, showing that (i) phenological variation among individuals strongly decreases under short day conditions (Figs. 2b and 3b) and (ii) genetic differences in day-length requirements are the single most important factor explaining variation in budburst times (Fig. 4c, d).

**Figure 4.**
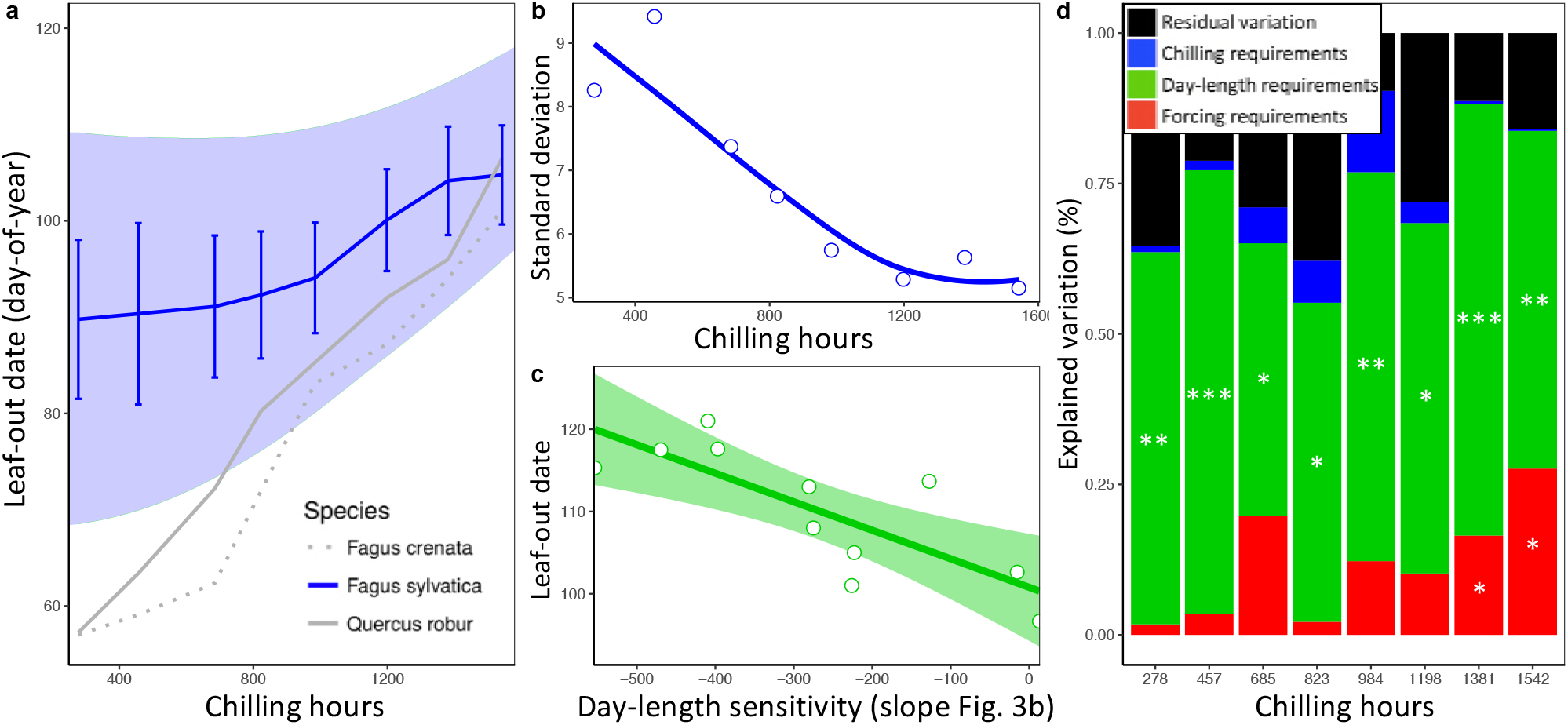
Loss of phenological synchrony with climate warming is explained by contrasting day-length sensitivities in *Fagus sylvatica*. **a, b**, Experiment 2. **a**, Leaf-out dates of *Fagus sylvatica* (blue), *Fagus crenata* (dotted grey), and *Ouercus robur* (grey) under varying winter lengths (chilling hours = sum of hours from 1 November until leaf-out with an average temperature between 0°C and 5°C). Bars show the standard deviation of average leaf-out dates among 11 *F. sylvatica* individuals. The shaded area shows the difference between the leaf-out date of the first flushing twig of the first individual and the last twig of the last individual to leaf-out, using a LOESS smoothing function. For *F. crenata* and *O. robur*, we investigated one individual each and therefore do not report inter-individual variation. **b**, Standard deviation of leaf-out dates among 11 *F. sylvatica* individuals at different winter lengths (chilling levels) and natural day length. **c**, The effect of individual day-length sensitivity on the timing of leaf unfolding when twigs were collected on 10 December 2015. Note the reversed x-axis scale, i.e., smaller values indicate higher day-length sensitivity. **d**, Variables explaining the sequence of leaf-out dates of 11 *F. sylvatica* individuals at eight different chilling levels. The percentage of leaf-out variation (derived from the ANOVA sums of squares) that can be explained by individual forcing requirements (red), day-length requirements (green), chilling requirements (blue), and the remaining residuals, i.e., unexplained variation (black). \**P* <0.05; \*\**P* <0.01; \*\*\**P* <0.001.

This insight explains why, especially in *Fagus sylvatica*, in which day length has the most pronounced effect on spring phenology (*10,11*), LOS is strongly affected by preseason temperatures (Fig. 1c). By contrast, in day-length insensitive species, such as silver birch *Betula pendula* and Norway spruce *Picea abies* (*11*), preseason warming has a smaller (but still significant) effect on LOS, suggesting that heritable differences in day-length sensitivity are a major driver of within-population phenological variation. In our common garden data, the standard deviation of inter-individual leaf-out times increased by 0.09 ± 0.02 (mean ± CI) days per decrease in one chilling day, and the standard deviation of inter-individual forcing requirements increased by 0.23 ± 0.06 degree-days per decrease in one chilling day (lower panel Fig. 2b), indicating that individual differences in the sensitivity to winter chilling also contribute to the observed loss of phenological synchrony under climate warming.

What biological consequences can be expected from less synchronized leaf-out and flowering of the individuals of a species? With regard to vegetative development, precocious leaf unfolding under warm springs increases the risk of late frost damage (*16–18*), but also potential carbon gain due to earlier photosynthetic activity (*19*). This risk-return trade-off will affect selection on suitable genotypes under future conditions, and the increasing spread of leaf-out should increase the selective importance of spring phenology. Whether opportunistic phenological strategies (relying on temperature as the main trigger) or conservative strategies (relying on day length and/or winter chilling as a buffer against highly variable spring temperatures) will be favored in the future will be region-specific, depending on the relative advancement rates of spring warming and late frost events. In continental regions, where the advent of spring is relatively invariable (low late frost risk), phenological strategies reliant on temperature should be favored (*20*).

With regard to flowering, decreased synchrony among individuals, as already strongly evident in *Alnus glutinosa* (Fig. 1f), should lead to reduced inter-individual pollen transfer. Strong divergence in flowering times among individuals also might lead to assortative mating (depending on incompatibility systems), possibly promoting local adaptation (*21–23*) and should act as a buffer against climate change-induced phenological mismatch between plants and leaf-feeding or pollen-collecting insects (*24*). Rapid adaptive responses, for instance a filtering out of extreme phenotypes through increased mortality or reduced reproduction, might counteract warming-induced losses of inter-individual synchrony. Such selection of the standing variation can occur very rapidly, at least in herbaceous plants (*25*).

While our results show that climate warming causes a loss of phenological synchrony among the individuals of a population, a study of leaf-out along elevational gradients in four European tree species, between 1960-2016, revealed that leaf-out times at higher and lower elevations are today compressed into a shorter time window compared to 58 years ago (*26*).

These findings do not contradict those of the present study because populations growing at high elevations were able to advance their phenology more than those at lower elevations for which chilling and/or day-length requirements are no longer fulfilled (Fig. S8). As a result, the leaf-out times of high- and low-elevation populations are converging (*26*). At the same time, however, differences in day-length sensitivity (as well as chilling and temperature sensitivity) among the individuals at any one elevation under climate warming are resulting in diverging flowering and leaf-out times.

The overall prediction from the present findings is that human-caused climate warming is leading to plant phenologies that are more heterogeneous within populations and more uniform among populations (over altitude or latitude). The rapid loss of reproductive and vegetative synchrony in European plants also predicts changes in their gene flow and trophic interactions, although community-wide consequences are presently unknown.

## Conclusion

The synchrony of developmental stages among organisms is a critical aspect of ecosystem functioning. Here, based on massive ground observations and climate-manipulation experiments, we show that global warming is altering within-population synchrony of leaf-out and flowering dates in temperate plants, with warmer temperatures reducing inter-individual synchrony by up to 55%. Experiments suggest that individual genetic differences in the sensitivity to day-length and/or winter chilling underlie the loss of synchrony, and future climate warming is expected to further strengthen this trend. These results predict consequences for gene flow and trophic interactions, but also emphasize the importance of adaptation when forecasting future plant growth and productivity.

## Materials and Methods

### Analysis of leaf-out and flowering synchrony (LOS and FLS) using the PEP database

#### Data sets

*In situ* phenological observations were obtained from the Pan European Phenology network (http://www.pep725.eu/), which provides open-access European phenological data. Leaf-out dates were analyzed for 9 species, flowering dates for 15. Data from Germany, Austria, and Switzerland were used for the analysis. For the angiosperm woody species, leaf-out was defined as the date when unfolded leaves, pushed out all the way to the petiole, were visible on the respective individual (BBCH 11, Biologische Bundesanstalt, Bundessortenamt und Chemische Industrie). For the two conifers *Larix decidua* and *Picea abies* leaf-out was defined as the date when the first needles started to separate (“mouse-ear stage”; BBCH 10). Flowering was defined as the date of beginning of flowering (BBCH 60). We removed (i) individuals, for which the standard deviation of phenological observations across years was higher than 25 and (ii) leaf-out and flowering dates that deviated from an individual’s median more than 3 times the median absolute deviation (moderately conservative threshold) (*26*).

#### Analysis

To test for an effect of spring temperature on inter-individual leaf-out synchrony (LOS) and flowering synchrony (FLS), we divided the study area into pixels of one degree resolution (~110 x 85 km), an area that can reasonably be considered as reflecting populations, at least for wind-pollinated woody species (see discussion on herbs in the main text). To allow for within-pixel comparisons of LOS and FLS between years, data from the same individuals had to be used each year. To achieve this, we kept only pixels for which there were at least three individuals with data for the same 15 years. For each pixel, we deleted all (i) individuals growing at altitudes that deviated by >200 m from the average altitude of all individuals within the pixel, and (ii) years that had less than 90% plant-coverage, i.e., data from at least 90% of the individuals within the pixel had to be available for the respective year, otherwise the year was excluded from the analysis. This data cleaning left us with a total of 12,536 individuals, 317,672 phenological observations (individuals x year), and a median time-series length of 25 years (minimally 15 years, maximally 48 years). The number of individuals within pixels (per species and phenological stage) ranged between 3 and 53 (median = 12). See Figs. S1b and S2b for information on the number of pixels used per species.

For each year and species, LOS and FLS within pixels were then calculated as the standard deviation of leaf-out or flowering dates. Additionally, we calculated the standard deviation of forcing requirements among individuals (subsequently referred to as LOS-DD [leaf-out synchrony degree-days] and FLS-DD [flowering synchrony degree-days]) to test if greater phenological variation among individuals can be explained by increasing variation in forcing requirements. Individual forcing requirements until leaf-out were calculated as the sum of degree-days (DD) from 1 January until leaf-out or flowering using 5°C as base temperature (e.g., ref. 27):

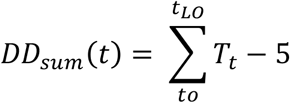

where *DD_sum_* is the accumulated degree days until leaf unfolding, *t_LO_* is the day of leaf unfolding, *T_t_* is the mean daily temperature on day *t*, and *t_0_* is the start date for forcing accumulation, which was fixed at 1 January. For each year and species, LOS-DD and FLS-DD within pixels were then calculated as the standard deviation of forcing requirements until leaf-out or flowering dates.

The daily mean air temperature at each site was derived from a gridded climatic data set of daily mean temperature at 0.5° spatial resolution (approximately 50 km, ERA-WATCH) (*28*). For each year, preseason temperature within pixels was defined as the average temperature during the 60 days prior to the average leaf unfolding or flowering date within the respective pixel, which is the period for which the correlation coefficient between phenological event and temperature is highest (*29*).

To test if shortened photoperiods and/or reduced winter chilling explain the decrease in phenological synchrony under warmer preseasons, for each year, pixel, and species, we calculated the average chilling hours until leaf-out or flowering and the average photoperiod (PP) at the date when the average forcing requirements until leaf-out or flowering were fulfilled. Chilling hours were calculated on basis of 6-hourly temperature data (CRU-NCEP, spatial resolution of 0.5°; https://crudata.uea.ac.uk/cru/data/ncep/), as the sum of hours from 1 November until leaf-out/flowering with an average temperature between 0°C and 5°C (e.g., ref 29):

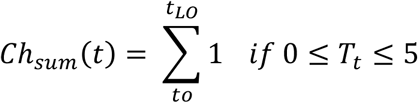

where *Ch_sum_* is the sum of chilling hours until leaf unfolding, *t_LO_* is the day of leaf unfolding, *T_t_* is the hourly mean temperature on hour *t*, and *t_0_* is the start date for chilling accumulation, which was fixed at 1 November in the year before leaf unfolding.

PP was calculated as a function of latitude and DOY (*30*):

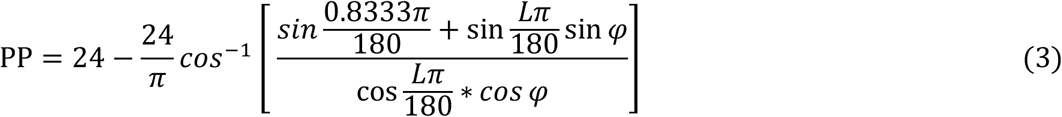

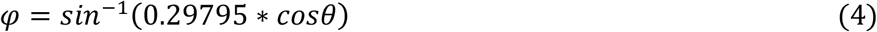

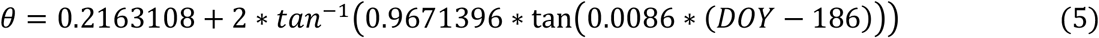

where *L* is the latitude of the phenological site.

#### Statistical analyses

Within each pixel we applied linear models to test for an effect of preseason temperature, photoperiod, and winter chilling on phenological synchrony (LOS, LOS-DD, FLS and FLS-DD). We then determined the frequency distributions for the correlation coefficients between phenological synchrony and preseason temperature across all species and sites. For each species, we applied *t*-tests to detect whether the average of all correlation coefficients obtained for each pixel differs from zero. To model changes in the distribution of within-population leaf-out and flowering dates (means and standard deviations) in response to temperature, we applied mixed-effects models using average leaf-out / flowering dates or LOS / FLS as response variables, preseason temperature as explanatory variable, and site as a random effect to control for the use of different sites in the model.

To test for the relative effects of preseason temperature on (i) inter-individual variation in leaf-out/flowering date (LOS / FLS) and (ii) inter-individual variation in forcing requirements until leaf-out/flowering (LOS-DD / FLS-DD) we applied hierarchical Bayesian models. To test for the effects of winter chilling and day-length on phenological synchrony, we applied hierarchical Bayesian models including both winter chilling until leaf-out and day length at the date when the average forcing requirements until leaf-out or flowering were fulfilled as predictor variables. The use of a Bayesian framework allowed us to fit slope parameters across traits simultaneously without concerns of multiple testing or *P*-value correction. All models included random effects for (i) species (to address within-species rather than between species phenological synchrony) and (ii) pixels (to address within-population rather than between-population phenological synchrony). To allow for direct effect size comparisons, all continuous variables were standardized by subtracting their mean and dividing by 2 SD before analysis (*31*). The resulting posterior distributions are a direct statement of the probability of our hypothesized relationships. Effective posterior means ± 95% confidence intervals are shown in Fig. 2.

To parameterize our models, we used the JAGS implementation (*32*) of Markov chain Monte Carlo methods in the R package R2JAGS (*33*). We ran three parallel MCMC chains for 200,000 iterations with a 50,000-iteration burn-in and evaluated model convergence with the Gelman and Rubin (*34*) statistic. Noninformative priors were specified for all parameter distributions, including normal priors for *α* and *β* coefficients (fixed effects; mean = 0; variance = 1,000), and uniform priors between 0 and 100 for the variance of the random intercept effects, based on de Villemereuil and colleagues (*35*). All statistical analyses relied on R 3.2.2 (*36*).

### Analysis of leaf-out synchrony (LOS) using common garden data from 2013–2018

Between 2013 and 2018 we observed the leaf-out dates of 209 individuals in 59 temperate woody species (minimally 3 individuals per species) in the Munich Botanical garden (see Supplementary Materials Table 1 for a list of species). An individual was scored as having leafed out when at least three branches had unfolded leaves pushed out all the way to the petiole (*37*). To test whether the trends observed in the PEP analysis are consistent with our common garden data, the same parameters (LOS, LOS-DD, preseason temperature, winter chilling, and photoperiod) were calculated as described above (*Analysis of leaf-out and flowering synchrony (LOS and FLS) using the PEP database*). We then applied hierarchical Bayesian models including species random effects (see paragraph above) to test for the effects of preseason temperature, winter chilling, and day-length on LOS and LOS-DD.

### Twig cutting experiments and phenological scoring

To study the extent of intraspecific variation in leaf-out strategy (within-species variation in photoperiod, chilling, and forcing requirements) and its implications under climate warming, we conducted twig-cutting experiments on mature *Fagus sylvatica* individuals grown in the vicinity of Munich. Experiments have demonstrated that twig cuttings precisely mirror the phenological behavior of their donor plants and therefore are adequate proxies for inferring phenological responses of adult trees to climatic changes (*27,38*). We used twigs approximately 50 cm in length, and immediately after cutting, we disinfected the cut section with sodium hypochlorite solution (200 ppm active chlorine), cut the twigs a second time, and then placed them in 0.5 l glass bottles filled with 0.4 l cool tap water enriched with the broad-spectrum antibiotics gentamicin sulfate (40 μg l^−1^; Sigma-Aldrich, Germany) (*11,27*). We then transferred the cut twigs to climate chambers and kept them under short (8 h), intermediate (12 h), or long day (16 h) conditions (see Experiment 1 below), or natural day length (Experiment 2 below). Temperatures in the climate chambers were held at 12°C during the night and 20°C during the day, with an average daily temperature of 16°C to simulate forcing temperatures. Illuminance in the chambers was about 8 klux (~100 μmol s^−1^ m^−2^). Relative air humidity was held between 40% and 60%. To account for within-individual variation, we used 10 replicate twigs per individual treatment and monitored bud development every second day. For each individual and treatment, we then calculated the mean leaf-out date out of the first eight twigs that leafed out. A twig was scored as having leafed out when three buds had unfolded leaves pushed out all the way to the petiole (*37*). Forcing requirements until leaf-out were calculated as the sum of degree-days [outside of and in climate chambers] from 10 December (1^st^ collection date) until leaf-out using 5°C as base temperature (e.g., ref. 27). Chilling hours were calculated as the sum of hours from 1 November until leaf-out with an average temperature between 0°C and 5°C.

### Experiment 1: Differences in photoperiod sensitivity among Fagus sylvatica individuals

In winter 2014/2015, twigs of 11 individuals (10 replicate twigs per individual and treatment) of *Fagus sylvatica* were collected at three dates during winter (22 Dec 2014, 6 Feb 2015, and 21 Mar 2015) and brought into climate chambers. Additionally, we collected twigs from one individual each of *Fagus crenata* and *Quercus robur*. Temperatures in the chambers ranged from 12°C during night to 20°C during day, with an average daily temperature of 16°C. Day length in the chambers was set to 8h, 12h, or 16h.

Individual photoperiod sensitivity was defined as the slope of the function between day-length treatment and accumulated degree days (>5°C) until leaf-out (twigs were collected on 21 March; see Fig. 3b). The steeper the slope, the stronger the effect of photoperiod on the amount of warming required for leaf-out. A flat slope indicates that photoperiod has no effect on the timing of leaf-out.

Individual chilling sensitivity was defined as the slope of the function between chilling treatment (collection date) and accumulated degree days (>5°C) until leaf-out when twigs were kept under constant 16-h day length (see Fig. 3c). The steeper the slope, the stronger the effect of chilling on the amount of warming required for leaf-out.

Individual forcing requirement was defined as the accumulated degree days (>5°C) until leaf-out under long chilling (21 March collection) and constant 16-h day length (see Fig. 3a). Under such conditions, chilling requirements and photoperiod requirements should be largely met, and thus the remaining variation in leaf-out dates should be largely attributable to differences in forcing (warming) requirements.

### Experiment 2: Different reactions to climate warming among Fagus sylvatica individuals

In winter 2015/2016, twigs from the same 11 individuals were harvested every two weeks (from 10 December until 21 March) and kept under the same temperature conditions applied in experiment 1 (12°C during night to 20°C during day), with natural day length. This allowed us to test if those individuals with no/little photoperiod sensitivity would advance their leaf-out more under short winter conditions than photoperiod-sensitive individuals, and to determine the relative effect of individual variation in photoperiod requirements, chilling requirements and forcing requirements on leaf-out variation under different winter/spring conditions (Fig. 4). Within-species leaf-out synchrony (LOS) was calculated as the standard deviation of individual leaf-out dates. To analyze which leaf-out cues (photoperiod, chilling, and forcing requirements) best explain leaf-out variation among individuals, we applied a multivariate linear model, including individual forcing, photoperiod, and chilling requirements (as inferred from experiment 1) as explanatory variables. To express the total variation in leaf-out dates that can be attributed to each trait, we used ANOVA sums of squares (see Fig. 4d).

To infer which percentage of the variation in leaf-out dates is due to treatment effects, between-individual variation, or within-individual variation, we calculated variance components by applying a random-effects-only model including treatments and individuals as random effects (individuals nested within treatments). Results show that of the total leaf-out variation among twigs, 52% can be explained by between-individual variation, 33% by treatments, and only 15% by within-individual variation (Supplementary Fig. S9).

## Acknowledgements

We thank D. März and V. Sebald for help with the experiments and R. Ricklefs for comments on the manuscript. This work benefitted from the sharing of expertise within the DFG priority program SPP 1991.

## Statement of authorship

CMZ designed the study, performed the experiments and analyzed the data. LM contributed to the analyses. CMZ and SSR wrote the manuscript.

## Supplementary Material

**Figure S1.**
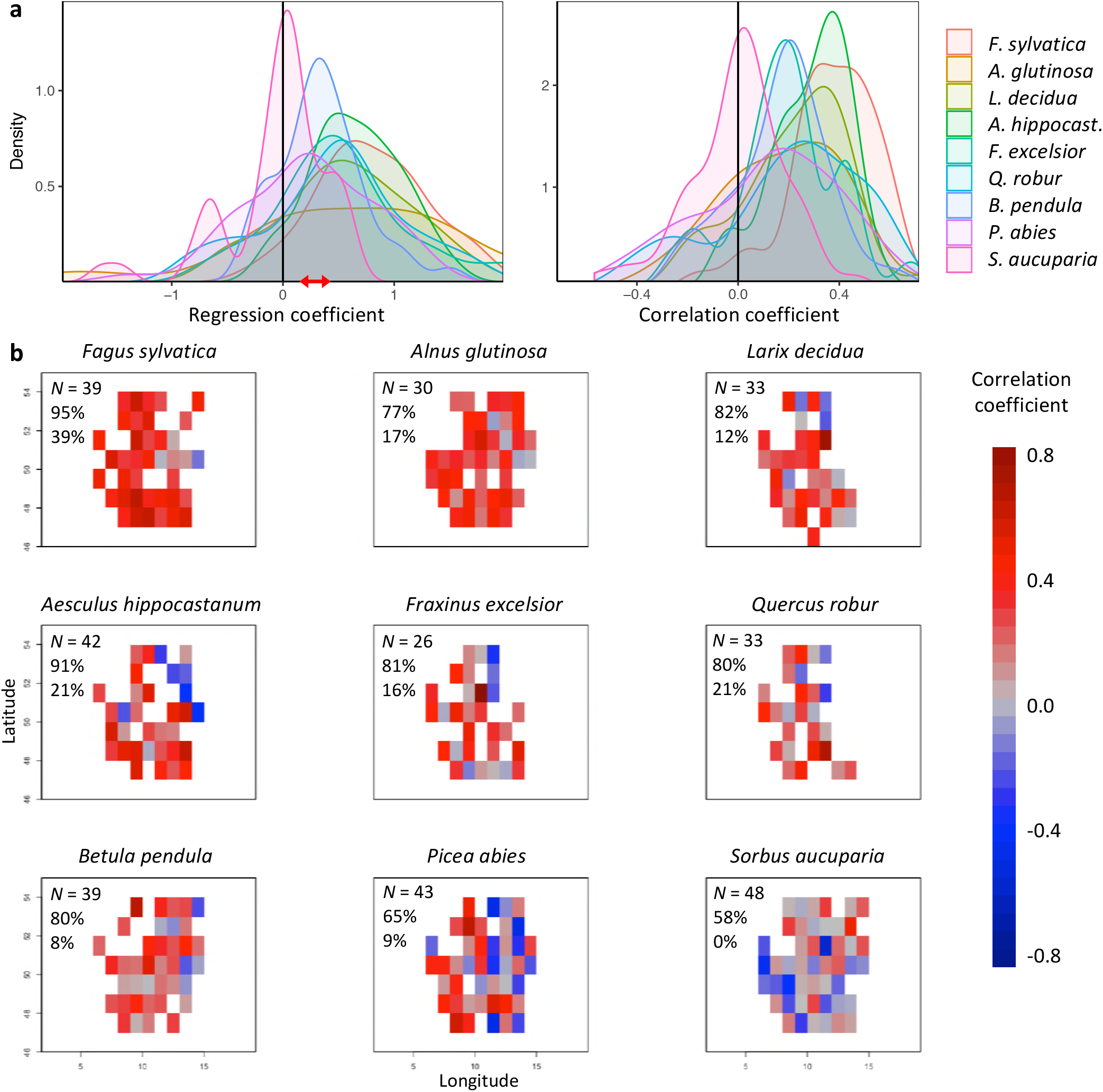
Effects of preseason temperature on inter-individual leaf-out synchrony (LOS), using PEP data. **a**, Density plots of regression (left) and Pearson correlation coefficients (right) between LOS and preseason temperature for nine species. Regression coefficients show the change in LOS per each degree increase in preseason temperature. The red arrow indicates the range of regression coefficients obtained when simulating spring phenology with a degree-day model (see Extended Data Fig. 5). **b**, Maps showing the regression coefficients for the effect of temperature on LOS at each site (colour coding according to correlation coefficients). *N* = Number of sites (1° x 1° pixels) in which the relationship was analysed. Percentages are the proportion of positive correlations and significantly positive correlations, respectively.

**Figure S2.**
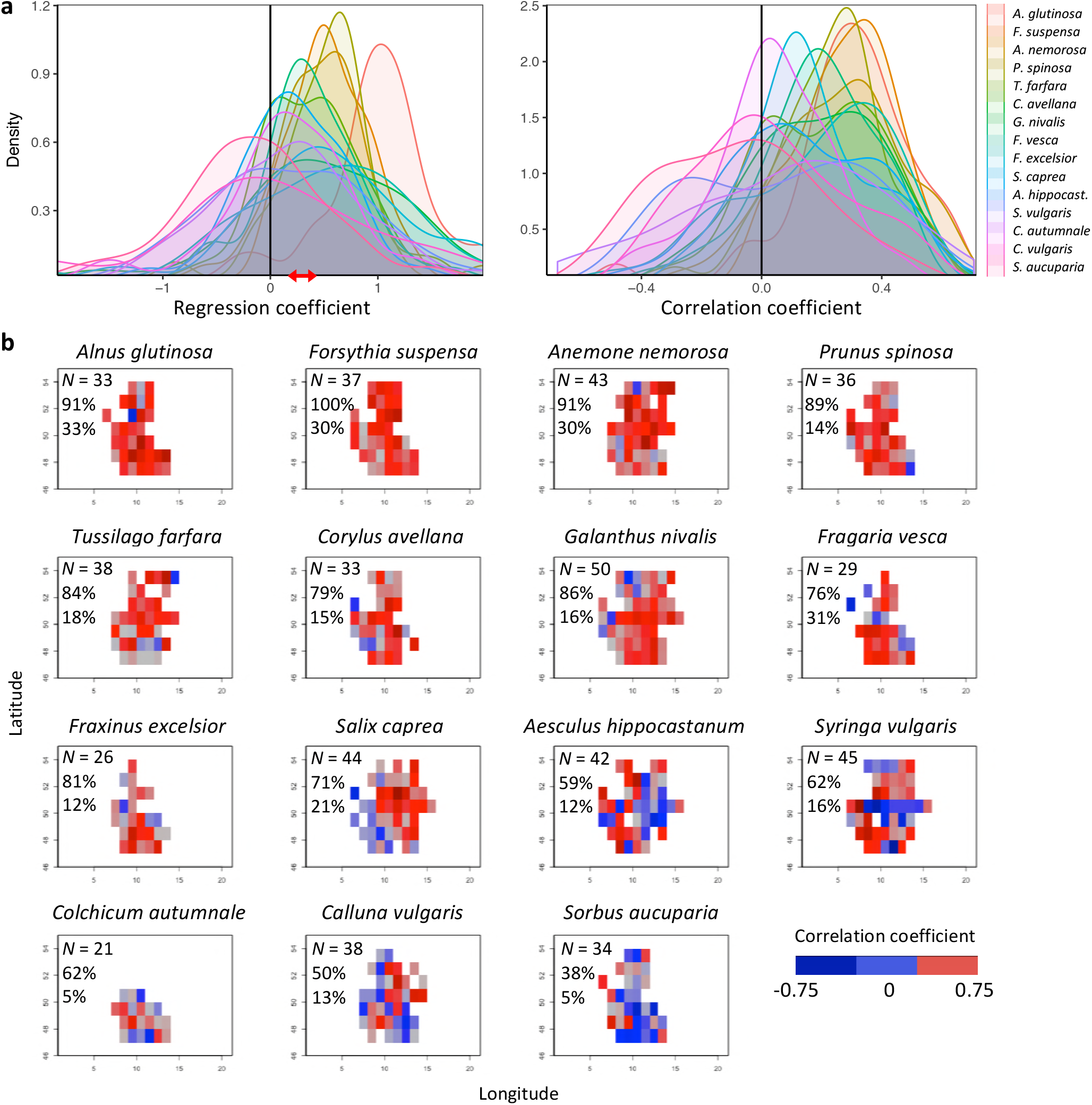
Effects of preseason temperature on inter-individual flowering synchrony (FLS), using PEP data. **a**, Density plots of regression (left) and Pearson correlation coefficients (right) between FLS and spring temperature for 15 species. Regression coefficients show the change in FLS per each degree increase in spring temperature. The red arrow indicates the range of regression coefficients obtained when simulating spring phenology with a degree-day model (see Extended Data Fig. 5). **b**, Maps showing the correlation coefficients for the effect of temperature on FLS at each site (colour coding according to correlation coefficients). *N* = Number of sites (1° x 1° areas) in which the relationship was analysed. Percentages are the proportion of positive correlations and significantly positive correlations, respectively.

**Figure S3.**
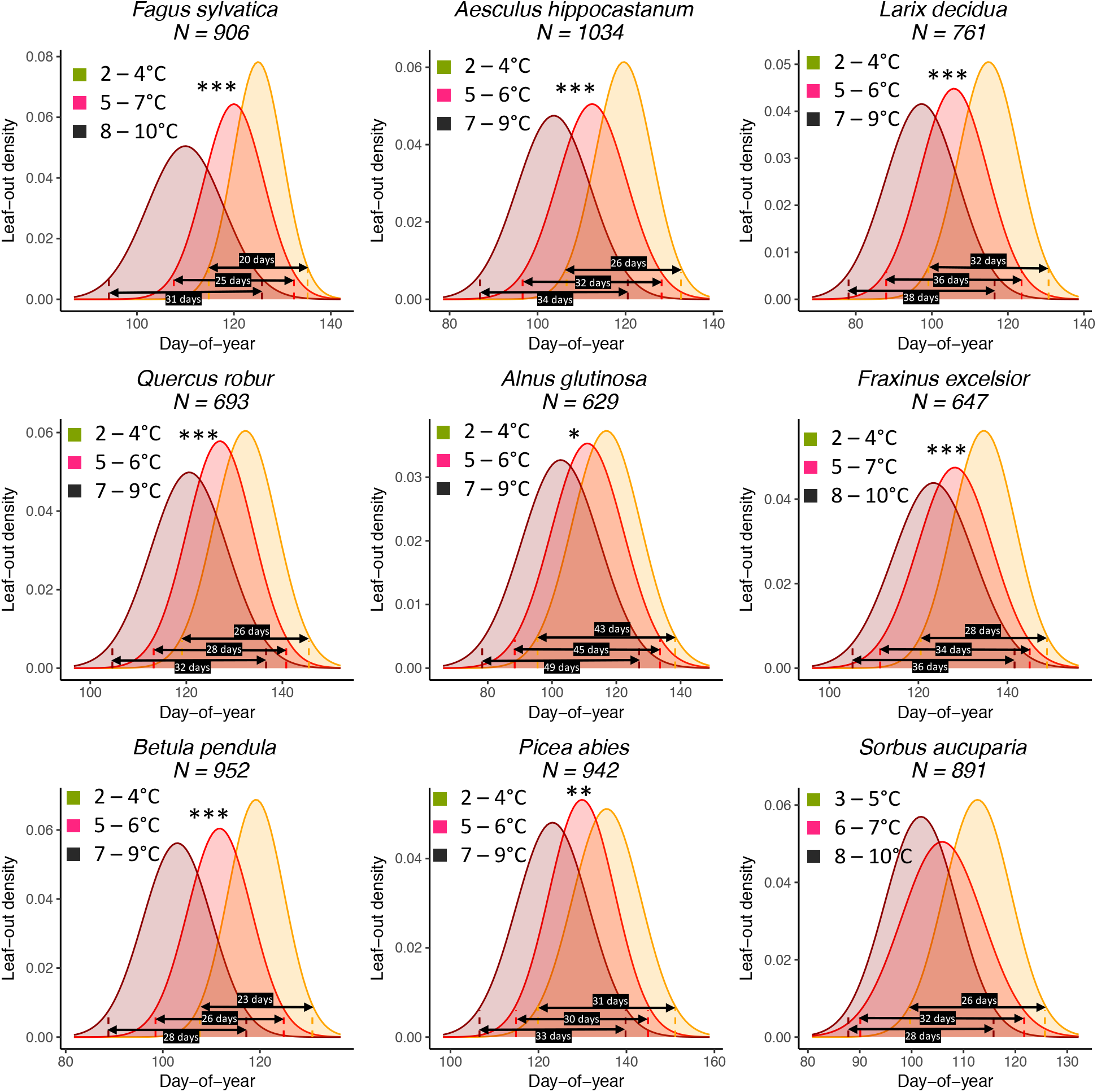
The effect of preseason temperature on inter-individual (within-population) leaf-out distributions. *N* = Number of available year x site (1° x 1° areas) combinations. To model the distributions (means and standard deviations), mixed-effects models were applied including site as a random effect. Stars indicate a significant positive effect of preseason temperature on LOS (\**P* <0.05; \*\**P* <0.01; \*\*\**P* <0.001). Black arrows show the period in which >95% of individuals leaf out (4 standard deviations), e.g., for *Fagus sylvatica*, in years with a cool preseason, 95% of individuals within a population leaf out within 20 days, whereas in years with a warm preseason this period is 31 days (55% longer).

**Figure S4.**
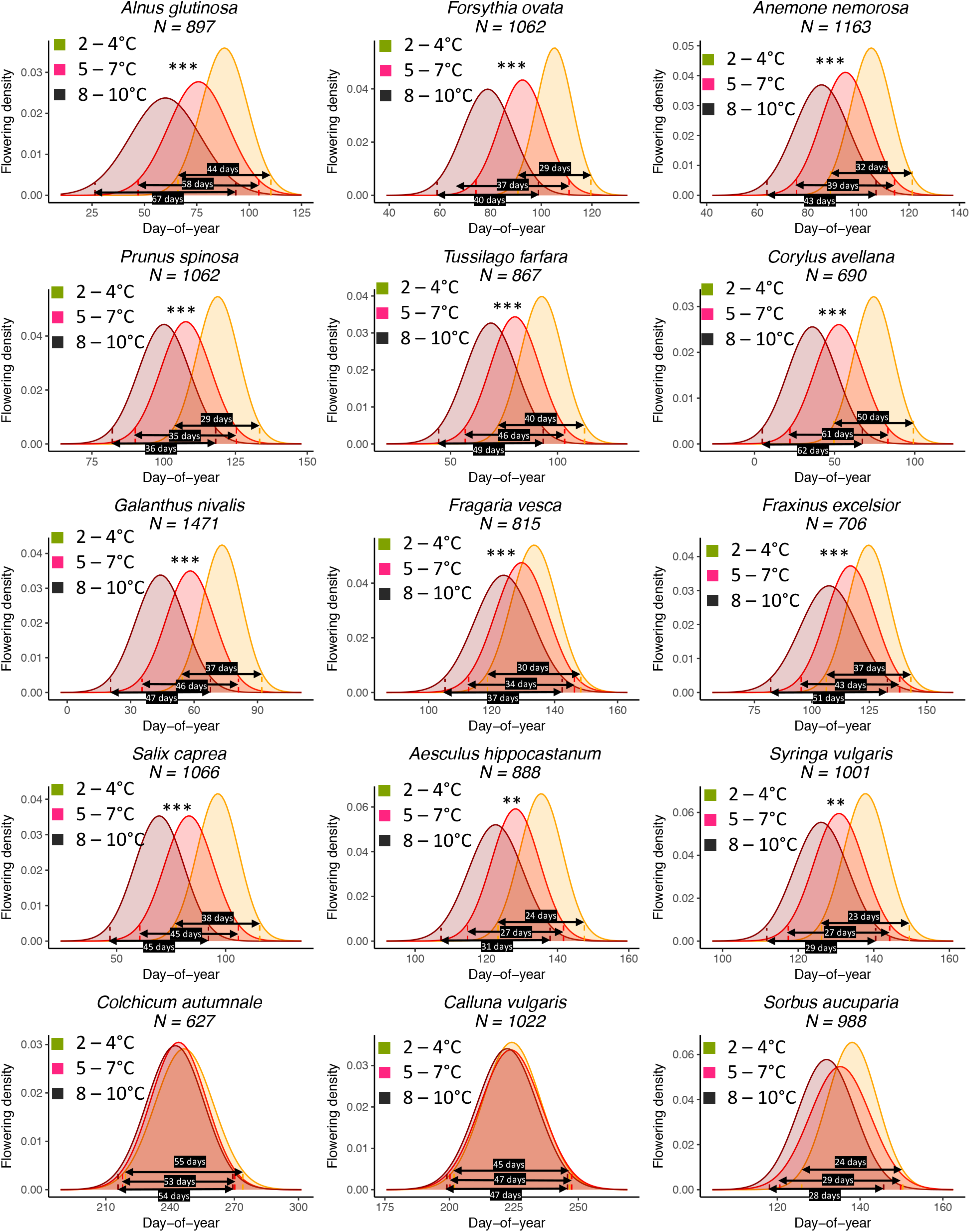
The effect of preseason temperature on inter-individual (within population) flowering distributions. *N* = Number of available year x site (1° x 1° areas) combinations. To model the distributions (means and standard deviations), mixed-effects models were applied including site as a random effect. Stars indicate a significant positive effect of preseason temperature on FLS (\**P* <0.05; \*\**P* <0.01; \*\*\**P* <0.001). Black arrows show the period in which >95% of individuals flower (4 standard deviations), e.g., for *Alnus glutinosa*, in years with a cool preseason, 95% of individuals within a population flower within 44 days, whereas in years with a warm preseason this period is 67 days (52% longer).

**Figure S5.**
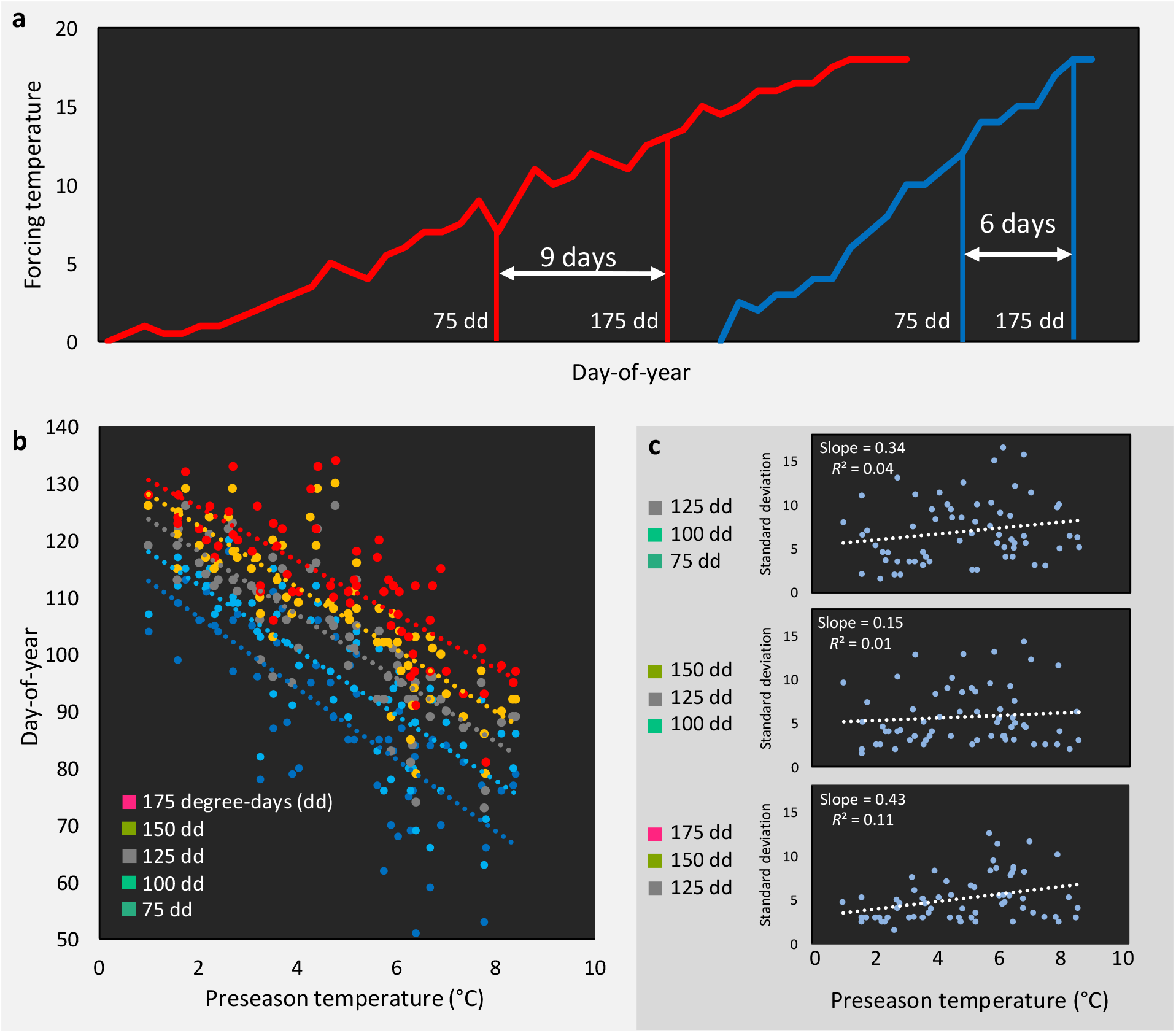
Does decreased LOS and FLS under climate warming result from a decrease in within-spring warming speed? **a**, Schematic representation of the hypothesized relationship between climate warming and within-spring temperature increase (see refs. 14,15): in a cold year (blue line) temperature rises late but fast, in a warm year (red line) temperature rises early but more slowly (flattening the temperature curve during spring). The result would be a less synchronized phenology in warm years, because forcing sums required for the occurrence of the respective phenological event are more spread out. This is illustrated by the date (day-of-year) at which 75 or 175 degree-days (cumulative daily temperature above 5°C starting 1 of January) accumulate in both cases (difference of 9 day in the warm year *vs*. 6 days in the cold year). **b**, The day of year when 75, 100, 125, 150, or 175 degree-days have accumulated, shown as response to mean preseason temperature (14 Feb until 15 April) in the respective year, using temperature data for 63 years (1955–2017) from Munich, Germany. **c**, The standard deviations of the dates (days of year) when (i) 75, 100, and 125 degree-days have accumulated (upper panel), (ii) 100, 125, and 150 degree-days have accumulated (middle panel), and (iii) 125, 150, and 175 degree-days have accumulated (middle panel) in response to preseason temperature.

**Figure S6.**
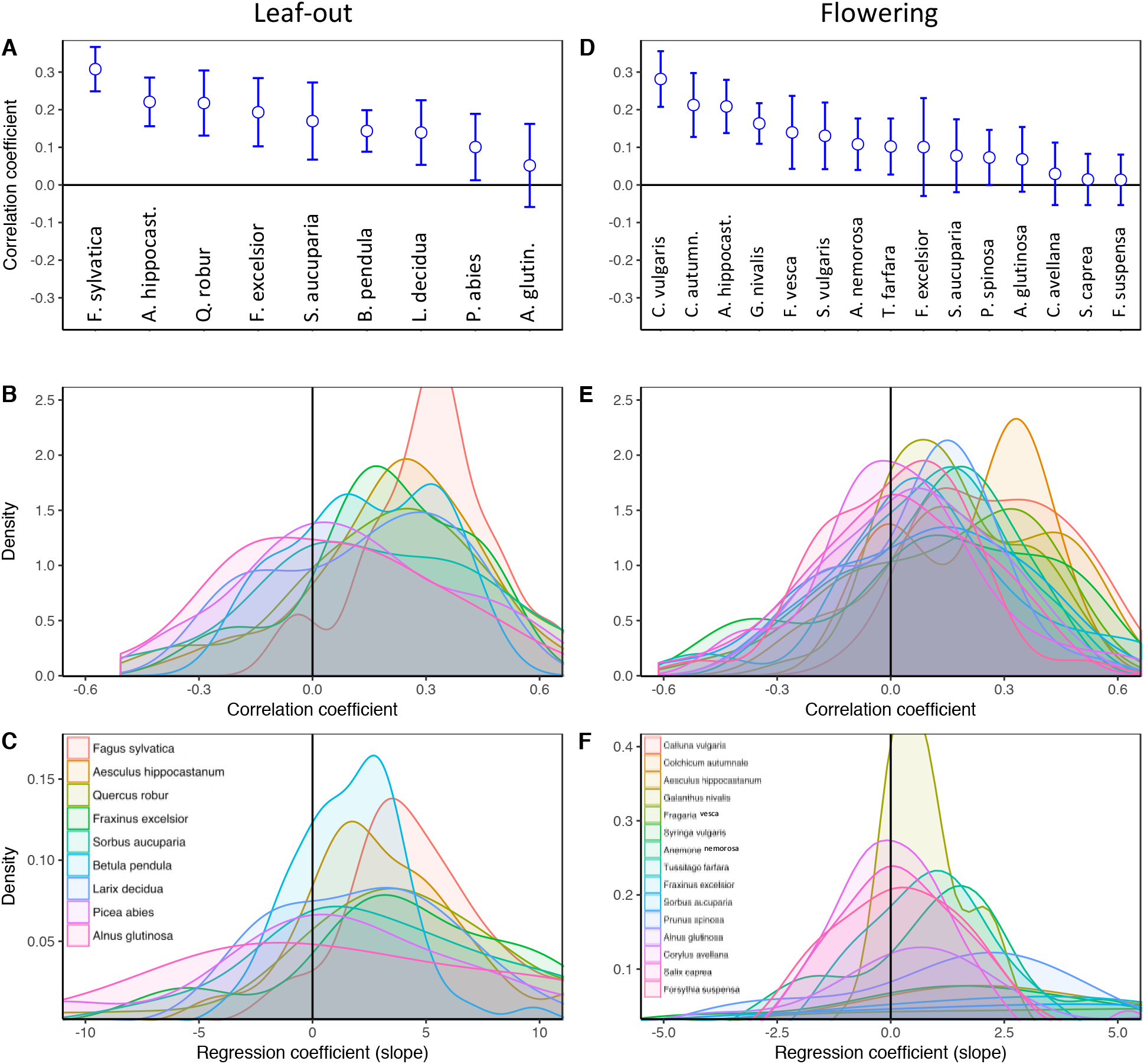
Greater variation of forcing requirements among individuals with increasing preseason temperatures. Effects of preseason temperature on inter-individual LOS-DD (**A–C**) and FLS-DD (**D–F**), using PEP data. **A**, Mean Pearson correlation coefficients (± 95% confidence intervals) for the effect of spring temperature on LOS-DD. See Figs. S1b and S2b for number of sites (1° x 1° areas) in which the relationship was analysed. **B, C**, Density plots of regression (**B**) and Pearson correlation coefficients (C) between LOS-DD and spring temperature for nine species. Regression coefficients show the change in LOS-DD per each degree increase in spring temperature. **D–F**, Same plots for the effect of spring temperature on FLS-DD. LOS-DD = Standard deviation of inter-individual forcing requirements until leaf-out; FLS-DD = Standard deviation of inter-individual forcing requirements until flowering.

**Figure S7.**
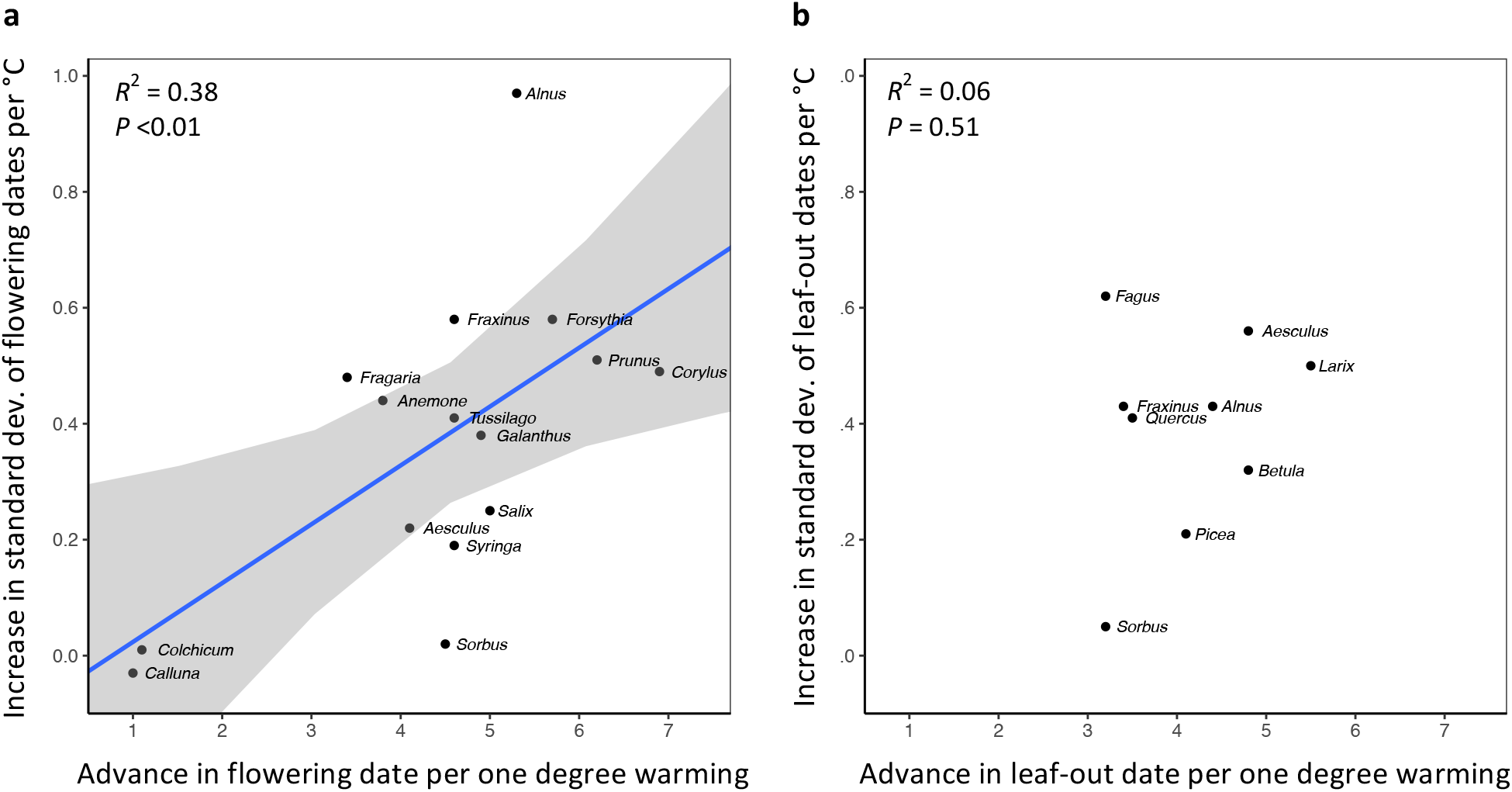
In species in which preseason temperature has little effect on the mean flowering date, preseason temperature also has little effect on FLS. **a**, Positive correlation between species’ mean temperature sensitivity of flowering date (days advance in flowering per one degree warming) and the mean temperature sensitivity of FLS (increase in the standard deviation of inter-individual flowering times per one degree warming). **b**, No correlation between species’ mean temperature sensitivity of leaf-out date (days advance in leaf-out per one degree warming) and the mean temperature sensitivity of LOS (increase in the standard deviation of inter-individual leaf-out times per one degree warming). The effects of preseason temperature on mean flowering date, mean leaf-out date, FLS, and LOS were inferred from mixed-effects models including site as a random effect. Species: *Alnus glutinosa, Aesculus hippocastanum, Anemone nemorosa, Betula pendula, Corylus avellana, Colchicum autumnale, Calluna vulgaris, Fraxinus excelsior, Forsythia suspense, Fagus sylvatica, Fragaria vesca, Galanthus nivalis, Larix decidua, Picea abies*, *Prunus spinosa*, *Quercus robur*, *Sorbus aucuparia*, *Salix caprea*, *Syringa vulgaris*, *Tussilago farfara*.

**Figure S8.**
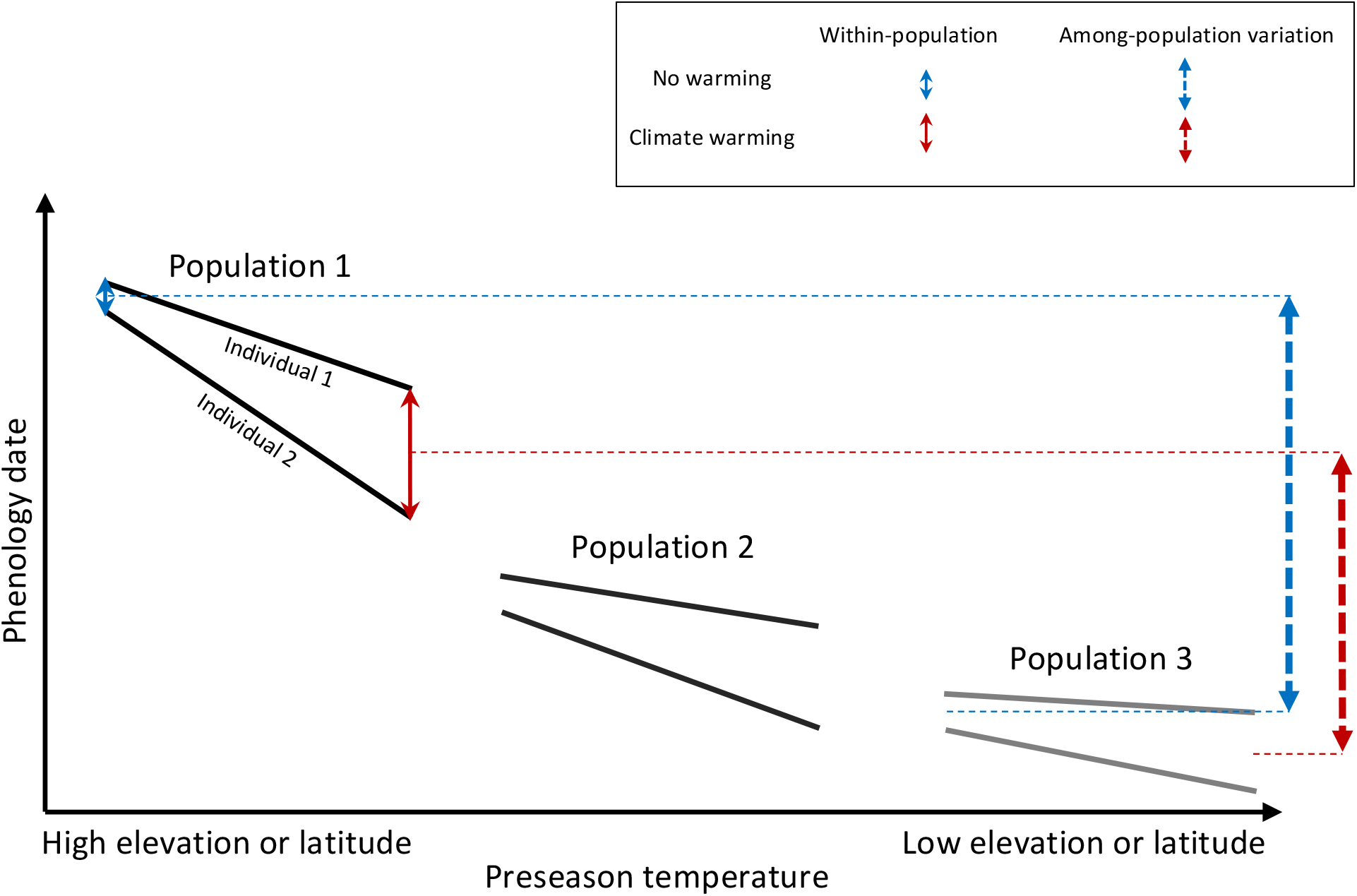
Schematic representation of within- and among-population phenological synchrony in response to climate warming. As demonstrated in this study, inter-individual synchrony within a population will decrease under warmer preseason temperatures because individuals differ in their sensitivity to temperature. Within-population variation under ambient or warmed preseason temperatures is illustrated by the solid blue and red arrows, respectively. By contrast, phenological synchrony among populations is expected to increase, given that populations in warm regions (Population 3) will advance their phenology less than populations in cold regions (Population 1). This is illustrated by the dashed blue and red arrows, showing that the difference in the average phenological date between Population 1 and 3 is smaller under warmer preseasons (red dashed arrow) than under ambient preseason temperatures (blue dashed arrow).

**Figure S9.**
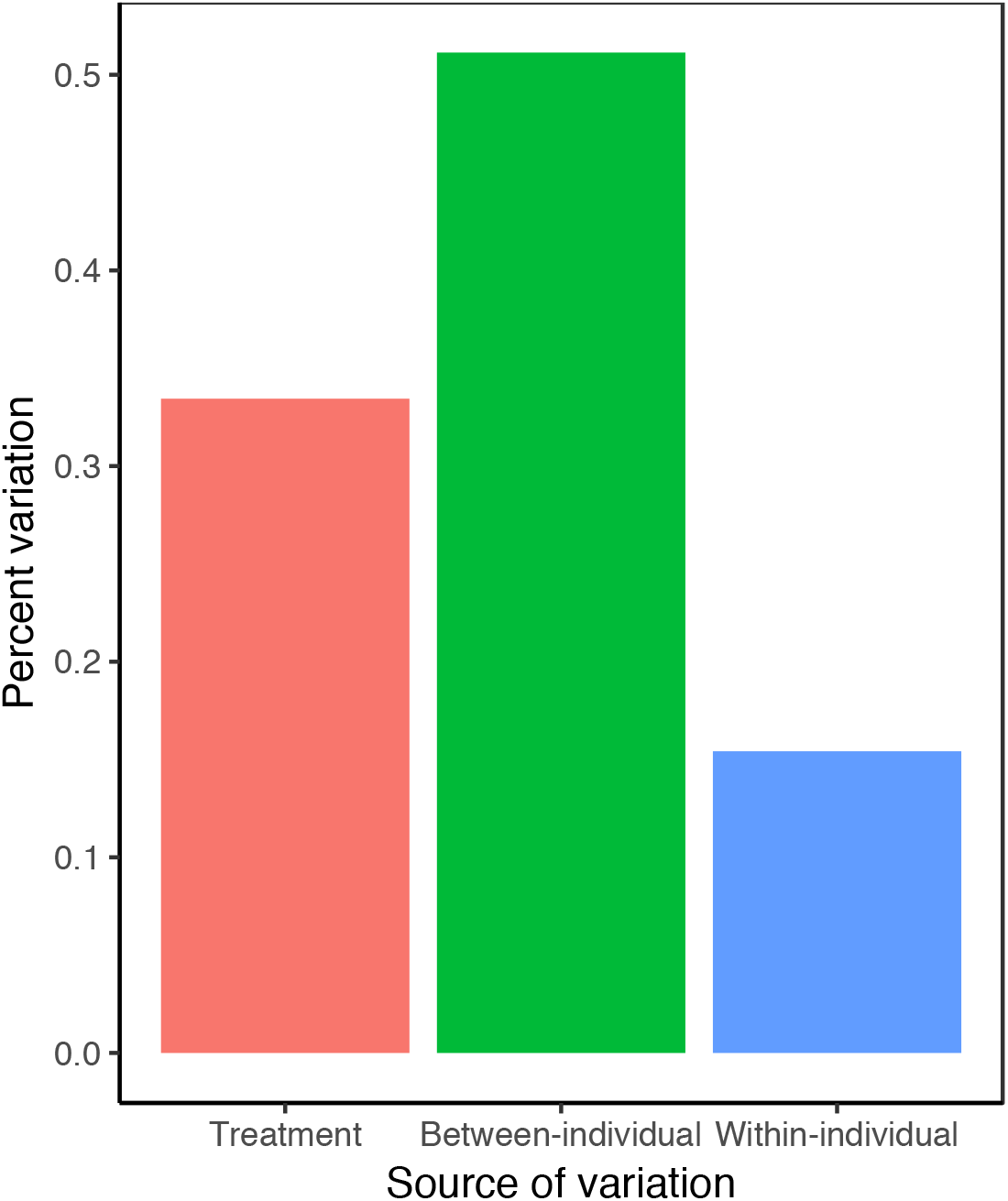
Percent variation in leaf-out dates attributable to treatment effects and between- and within-individual variation within treatments. Data from experiment 2 (see Methods). Variance components were inferred from random-effects-only models, including leaf-out date as the dependent variable and treatment and individuals as nested random effects.

